# INVESTIGATION OF ETHYLENE OXIDE GENOTOXICITY DOSE-RESPONSE TO INFORM CANCER RISK ASSESSMENT

**DOI:** 10.64898/2026.03.25.714257

**Authors:** B. Bhaskar Gollapudi, James E. Bus, Phillip Cassidy, Jeffrey T. Weinberg, Jeffrey C. Bemis, Dorothea. K. Torous, Stephen D. Dertinger, Kun Lu, Abby A. Li

## Abstract

Ethylene oxide (EtO) is primarily used as an intermediate in the manufacture of chemicals, with a minor use as a sterilant for medical equipment and food products. It is a direct-acting alkylating agent that reacts with cellular macromolecules, including proteins and DNA. EtO has been shown to induce tumors in rodents and humans. DNA reactivity has been the postulated mode of action (MOA) for its carcinogenicity. The current study has investigated the dose response for EtO-induced genetic damage to inform the biological plausibility of a dose-response model for cancer risk assessment. Male and female B6C3F1 mice were exposed to 0, 0.05, 0.1, 0.5, 1, 50, 100, or 200 ppm EtO by whole-body inhalation (6 hours/day for 28 days, 7 days/week). Mutagenicity was assessed by determining the frequency of mutant *Pig-a* phenotype in reticulocytes (RET) and mature red blood cells (RBC) on Day 28. Cytogenetic damage was evaluated by the erythrocyte micronucleus (MN) test in blood samples collected on Days 5 and 28. EtO is a relatively weak genotoxicant with treatment-related increases in *Pig-a* and MN frequencies being seen primarily at 200 ppm. The hockey-stick shaped dose response for genetic damage may be conservatively interpreted as being no more than a linear response with a single slope. Thus, a cancer risk assessment dose-response model consisting of a single linear slope throughout the exposure range is biologically plausible and consistent if EtO were acting through a mutagenic MoA for its carcinogenicity.

## INTRODUCTION

Ethylene oxide (EtO) is used as an intermediate in the manufacture of chemicals with approximately 1% of the manufactured EtO being used as a sterilant for medical equipment and food products. Human exposure to EtO occurs via natural endogenous and exogenous sources (Kirman et al., 2021; Kirman et al., 2025). Endogenously, EtO is produced via metabolism of ethylene produced by gut bacterial metabolism or secondary to systemically mediated oxidative metabolism. Exogenous sources of EtO exposures include occupational, emissions from industrial and sterilization facilities, cigarette smoke, combustion of fossil fuels, forest fires, volcanoes, and systemic metabolism of ethylene associated with ripening of fruits and vegetables.

EtO is an *S_N_*2-type alkylating agent that reacts with the nucleophilic sites on the DNA, RNA and proteins. Although several different 2’-hydroxyethyl adducts are induced in the DNA, with the *N7*-(2-hydroxylethyl)guanine (N7-HE-G) being the most abundant one (>80%; Li et al., 1992), the minor *O^6^*-hydroxylethylguanine (O^6^-HE-dG) adduct is generally believed to be responsible for the mutagenicity of EtO (Swenberg et al., 2011; Pottenger et al., 2019). EtO is genotoxic both *in vitro* and *in vivo*, inducing gene mutations and cytogenetic damage (EPA, 2016; Gollapudi et al., 2020). It requires relatively high exposure concentrations and longer exposure durations to elicit a genotoxic response in experimental *in vivo* systems (Donner et al., 2010; Gollapudi et al., 2020; Manjanatha et al., 2017; Parsons et al., 2013; Recio et al., 2004). Thus, its mutagenic potency is relatively weak.

EtO is carcinogenic at multiple sites in inhalation animal cancer bioassays (Lynch et al., 1984; NTP 1987; Snelling et al., 1984). Increased cancer risk was also reported in humans occupationally exposed to EtO. For example, positive exposure-response trends were found for males for lymphoid cancer mortality, and for females for breast cancer mortality, with statistically significant excesses for these two cancers at the highest cumulative exposures (Steenland et al. 2003, 2004). For one industrial cohort, however, no increases in lymphoid cancer mortality were reported in male workers in EtO production facilities (Swaen et al. 2009; Valdez-Flores et al. 2025). Workplace exposure limits ranged from 100 ppm in 1946, 50 ppm in 1957 and 1 ppm in 1984 (ACGIH, 2015; 8h TLVs). An average 4.6 ppm EtO occupational exposure has been reported for sterilizer facilities operating between 1979 and 1985 (Hornung et al. 1994). Higher EtO exposures have been estimated to be approximately 70-100 ppm in the 1940s in EtO production plants and sterilizer facilities (Bogen et al., 2019; Swaen et al., 2009).

The US EPA (2016) and the Texas Commission on Environmental Quality (TCEQ, 2020) both used epidemiology data from the same cohort of sterilizer workers exposed to EtO that was assembled by the National Institute for Occupational Safety and Health (NIOSH). Both agencies assumed a genotoxic mode of action (MOA) and assumed a “linear no-threshold” assumption for the extra cancer risk (US EPA, 2005). However, the cancer risk derived by the US EPA (1 in a million extra risk = 0.0001 ppb) is approximately three orders of magnitude lower than the equivalent 0.24 ppb TCEQ value. Two key factors contributed to this difference. First, the US EPA (2016) included both lymphoid cancer mortality and breast cancer incidence in the calculation of cancer risk, while the TCEQ considered lymphoid cancer mortality alone. Second, and of greater quantitative impact to the cancer risk estimate, the TCEQ employed a standard log-linear Cox proportional hazard (CPH) statistical model with a single-slope essentially linear response covering the entire range of occupational exposures (Figure 1). In contrast, the US EPA applied a 2-piece linear spline model with a steep initial linear CPH model at low exposures, followed by a shallower linear CPH model at higher exposures connected by a modeled point of inflection (or knot) at 1600 ppm-days (Figure 1). The TCEQ’s and US EPA’s models have comparable statistical fits to the individual hazard rate data that were modeled (TCEQ, 2020). The critical question is which model has greater biological plausibility for deriving EtO cancer risks. The US EPA Science Advisory Board (SAB) review of the US EPA risk assessment importantly observed that “any model that is to be considered reasonable for risk assessment must have a dose response form that is both biologically plausible and consistent with the observed data.” (EPA SAB, 2015). The aim of the investigation reported here is to examine the shape of the dose-response for EtO-induced genetic damage. An early key event in the genotoxic MOA for EtO-induced carcinogenesis is expected to involve the induction of a heritable genetic alteration (i.e., mutation). Thus, it was hypothesized that the dose-response genetic damage should inform whether either or both the standard log-linear CPH or 2-piece linear spline models are acceptable biologically plausible alternatives for deriving EtO cancer risks. Towards this end, the dose-response for the induction of *Pig-a* mutations and micronuclei (a reporter for cytogenetic damage) was investigated in the erythrocytes, along with other endpoints specific to mutagenic MOA, in mice exposed for 28 days via whole body inhalation to a 4000-fold range of EtO concentrations.

**Figure 1:**
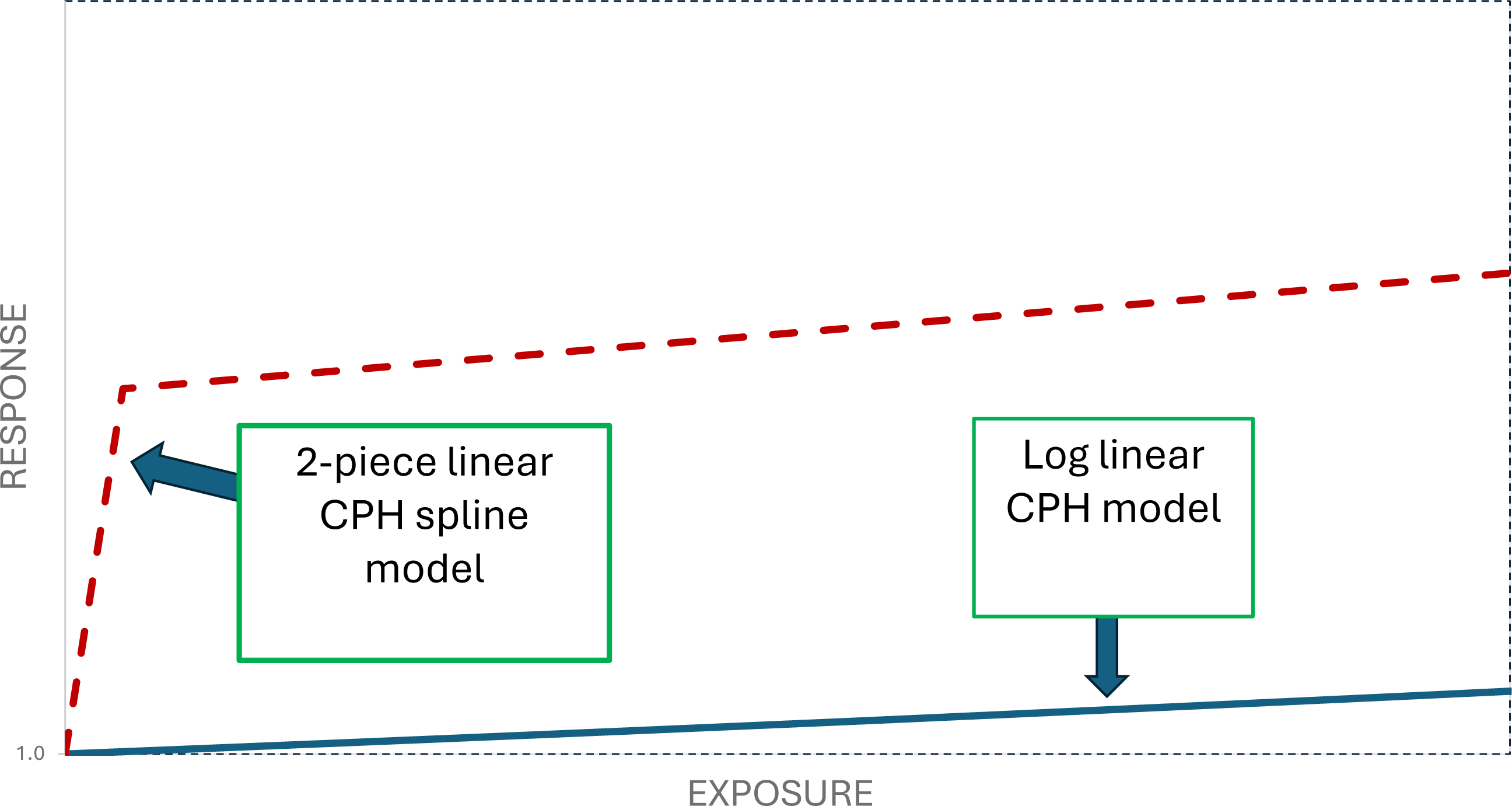
Graphical Representation of the 2-Piece Linear Spline and Log-Linear Cox-Proportional Hazard (CPH) Models used for Cancer Risk Assessment for Ethylene Oxide. Note: Response is graphed relative to the y-intercept of each type of hazard model; thus, they are comparable in terms of general shape but not along the y-axis (EPA, 2016 Figure 4-7; TCEQ, 2020 Appendix 6.3; Valdez-Flores et al. 2025)

## MATERIALS AND METHODS

### Animal Exposures and Tissue Sampling

A detailed description of the in-life procedures, including exposure generation and analytical monitoring, is presented in Liu et al. (2026a). The following is a brief overview of the methods. Ethylene oxide (CAS 75-21-8; purity: 99.99%) was provided by Balchem Corporation, Green Pond, SC, USA. The study protocol was reviewed and approved by the Institutional Animal Care and Use Committee of the laboratory performing the study (Charles River Laboratories, Ashland, OH, USA). Male and female B6C3F1/J mice (9-12 weeks old at study initiation) were purchased from The Jackson Laboratory, Bar Harbor, ME. Due to technical issues related to exposure generation and analysis of chamber concentrations in the two lowest exposure groups (i.e., 0.05 and 0.1 ppm; see below and Liu et al., 2026a) during the original execution of the study (hereafter referred to as Phase I), these were judged to be unreliable and needed to be repeated in a second phase (Phase II). The targeted concentrations evaluated were 0 (filtered air), 0.05, 0.1, 0.5, 1, 50, 100, and 200 ppm in Phase I and 0 (filtered air), 0.05, 0.1 and 50 ppm in Phase II. The 0 and 50 ppm exposures were repeated in Phase II to examine the reproducibility of the biological responses despite the differing exposure methodologies between the two phases. Filtered air and EtO were administered daily to mice as a 6-hour/day whole-body inhalation exposure for 28 consecutive days^1^. In Phase I, chemical and physical interference from animals in the chamber (e.g., a combination of odors, excess humidity from urine and/or feces, and animal dander) resulted in poor chromatography and high variability in EtO concentrations determined in the two lowest concentrations (0.05 and 0.1 ppm). Direct measurement of exposure concentrations in the 0.05 and 0.1 ppm chambers was terminated during the course of the study. Because of this, Phase I exposures, concentrations in the 0.05 and 0.1 ppm chambers were calculated and reported as nominal concentrations based on calibrated dilution air and EtO flow rates and analyzed concentrations of the 500 ppm intermediate dilution chamber. To alleviate this issue in Phase II, a serial chamber design was implemented so that EtO levels could be measured without animal interference.

Exposures were conducted using 1000-L stainless steel and glass whole-body exposure chambers. Animals were individually housed in batteries (consisting of linearly attached cages) and placed in the middle (Phases I and II) and lower (Phase I only) positions in the chamber. Food and water were withheld during exposure, but a water substitute (HydroGel® Recovery) was provided to each mouse. For the exposure of the filtered air group, supply air was used to achieve a similar humidity, airflow, and oxygen content to that in the test substance-exposed groups. In Phase I, an exposure atmosphere was generated by releasing neat test substance from generation bags and subsequent dilution to reach the target concentrations. In Phase II, an exposure atmosphere was generated for the 0.05, 0.1 ppm and 50 ppm groups by releasing neat test substance from generation bags into a 130-L mixing chamber, where it was mixed with supply air to reach the targeted concentrations. Exposure atmospheres were analyzed using a gas-phase injection gas chromatograph with a mass spectrometric detector (GC-MS) (Phase I, Groups 0, 0.05, 0.1, 0.5 and 1 ppm and Phase II, Group 0, 0.05, and 0.1 ppm), or a gas-phase injection gas chromatograph with a flame ionization detector (GC-FID) (Phase I, Groups 50, 100, and 200 ppm and Phase II, Group 50 ppm). Testing of exposure atmospheres took place at least three times and up to six times during each 6-hour exposure interval to ensure that the exposure concentration did not change over time. GC-MS analysis of exposure concentrations has never been used for EtO inhalation studies and is unique in general for inhalation testing but was highly discriminatory and provided the proper precision and accuracy necessary when dealing with concentrations as low as 0.05 ppm.

A group of animals dosed via oral gavage once daily for two consecutive days with cyclophosphamide monohydrate dissolved in deionized water (CP, CAS 6055-19-2, 25 mg/kg/day; Sigma-Aldrich, St. Louis, MO, USA) on Days 25 and 26 served as the positive control for micronucleus (MN) assay. Animals dosed once daily for three consecutive days with *N*-nitroso-*N*-ethyl urea dissolved in pH 6.0 phosphate-buffered saline (ENU ISOPAC®, CAS 759-73-9, 30 mg/kg/day; Sigma-Aldrich) on Days 1–3 via oral gavage served as positive controls for Day 5 MN and Day 28 *Pig-a* gene mutation assays.

Blood samples were collected at the end of exposure on Day 5 of the study for MN analysis (see below). The animals were euthanized immediately following the final exposure on Day 28, and samples of blood, liver, lung, mammary tissue (females), and bone marrow were collected from all surviving animals. Blood samples were used for assessing the frequencies of *Pig-a* mutations and MN (see below) and the formation of hemoglobin adducts (described in Liu et al., 2026a). Lung, liver, bone marrow and mammary tissues were used for the assessment of DNA adducts, and the results of these analyses were reported by Liu et al. (2026b). The adduct data served as a metric for the bioavailability as well as the exposure of the bone marrow tissue to the inhaled EtO.

### Flow Cytometric Analysis of Pig-a Mutant Frequencies

Blood samples from males and females were analyzed separately using MutaFlow® kits (Litron Laboratories) following the manufacturer’s instructions. Briefly, frozen whole blood samples were thawed and after centrifugation, anticoagulant was added to each. The entire volume of each was overlaid onto Lympholyte®-Mammal and the pellets resuspended and washed. An aliquot of each leukodepleted blood sample was added to a solution containing fluorescent antibodies to label wild-type erythrocytes (anti-CD24-PE), and fluorescent antibodies to label remaining platelets and leukocytes (anti-CD61-PE and anti-CD45-PE, respectively). The samples were incubated, centrifuged and the resulting pellets were resuspended, and an aliquot of Anti-PE MicroBeads was added to each. After incubation, samples were washed and resuspended. An aliquot of each Pre-Column sample was added to nucleic acid dye (with RNase) plus counting beads and incubated at ambient temperature, then placed on ice until analysis.

Each remaining labeled sample was added to an LS Column in a magnetic field. The Anti-PE MicroBeads retain most of the wild-type erythrocytes and contaminating platelets and leukocytes in the column, while the mutant-phenotype erythrocytes pass through into a centrifuge tube. After centrifugation, the cell pellets were resuspended, and an aliquot of nucleic acid dye plus counting beads was added to each. Each post-column sample was incubated at ambient temperature, then placed on ice until analysis. An Instrument Calibration Standard (ICS) was created by using a portion of unlabeled, leukodepleted blood and a portion of labeled, leukodepleted blood. Both portions were incubated in nucleic acid dye plus counting beads. Whereas true mutant cells are rare, the unlabeled blood mimics mutant cells, allowing for proper setting of PMT voltages and compensation, and establishing the demarcation line between mutant and wild-type cells. Each Pre-Column and Post-Column sample was analyzed by a BD FACSCalibur™ flow cytometer running CellQuest™ Pro software, version 5.2 (Becton Dickinson, San Jose, CA). Photomultiplier tubes collected the fluorescence emitted by each cell.

In general, a minimum of 100 million RBC and 3 million reticulocytes were evaluated from each animal to determine the mutant frequencies. In the few instances where this number could not be acquired, the sample size analyzed was always ≥40 million RBC and >2 million reticulocytes. The proportion of nucleic acid dye-positive RET among total erythrocytes (%RET) was calculated as an indication of bone marrow toxicity.

### Flow Cytometric Analysis of MN Frequencies

Blood samples from males and females were analyzed separately using MicroFlow® kits (Litron Laboratories) following the manufacturer’s instructions. Whole blood samples were diluted with an anticoagulant and fixed in ultracold fixative. All fixed blood samples were stored in a freezer (–90 °C to –80 °C) until analysis. Each fixed blood sample was washed with Hank’s Balanced Salt Solution (HBSS), and cells were isolated by centrifugation. An aliquot of each washed blood sample was added to a solution containing RNase, anti-CD71 FITC (an antibody to the transferrin receptor to label RETs), and anti-CD61-PE (an antibody to label platelets). The samples were incubated at 2 °C to 10 °C and then at ambient temperature. After incubation, the cells were kept at 2 °C to 10 °C until analysis. A propidium iodide (PI) solution was added to each sample to stain all DNA, including MN in the cells.

Methanol-fixed blood from mice infected with *Plasmodium berghei* and methanol-fixed blood from uninfected rats were used to configure the flow cytometer before analysis. Each blood sample was analyzed by flow cytometry on a BD FACSCalibur. Up to 20,000 reticulocytes (RETs; CD71+) were evaluated for the presence of MN. In addition, the number of normochromatic erythrocytes (NCE) and MN-NCE encountered in enumerating the desired number of RETs was also determined. The proportion of CD71 positive RET among total erythrocytes (%RET) was calculated as an indication of bone marrow toxicity.

### Statistical Analysis

All statistical analyses of data from the EtO groups were conducted using JMP Pro 18.1.1 (Macintosh). The sexes were analyzed separately, as were each time point and endpoint combination. Homogeneity of variance was assessed using Levene’s test. Box-Cox transformations were applied when needed to reduce heteroscedasticity. Initially, the two phases were evaluated using fixed-effects two-way ANOVA restricted to dose groups that were present in both phases to test for a Treatment x Phase interaction effect. A non-significant Treatment × Phase interaction was used to support pooling data across phases. Once pooling was justified, the full dataset (all dose levels from both Phases) was analyzed using a refined two-factor ANOVA that included Treatment Group and Phase as fixed effects and omitted the non-significant interaction term. Retaining Phase in this refined model ensured that treatment effects were appropriately adjusted for any underlying Phase differences, even when such differences were subtle or not statistically significant. For endpoints showing a significant Treatment effect in the pooled model, post-hoc Dunnett’s comparisons to the concurrent control were performed within JMP’s Fit Model platform so that all contrasts were based on least-squares means adjusted for Phase. This approach maintains internal consistency between the ANOVA model and the subsequent pairwise comparisons and yields treatment comparisons that accurately reflect the study’s two-phase structure. Statistical significance was inferred for pair-wise comparisons when the p value was <0.05 (one-sided) for *Pig-a*, MN-NCE and MN-PCE, 0.05 (2-sided) for %RET and 0.025 for trend tests. Given the positive control groups were included to ascertain the adequacy of the sample handling for flow cytometry, data on *Pig-a* and MN frequencies from these groups were not statistically analyzed but rather evaluated based on expert judgement.

## RESULTS

The current investigation aimed at elucidating the shape of the dose-response for EtO-induced erythrocyte *Pig-a* gene mutations and MN as reporters of bone marrow genotoxicity following whole body inhalation exposure. Blood from this study was used to investigate hemoglobin adducts as a biomarker for systemic EtO exposures, while various tissues (lung, liver, bone marrow and mammary tissue) were assessed for DNA adducts (Liu et al, 2026a, b). There was an exposure-dependent increase in the formation of both hemoglobin and the N7-HE-G DNA adducts at all concentrations of EtO tested. However, mutagenic O^6^-HE-dG adducts were only detected in mice exposed to ≥50 ppm EtO using a very sensitive technique with a detection limit of less than 1 adduct/cell. The N7-HE-G adduct data reported by Liu et al. (2026b) for the bone marrow is graphically presented in Figure 2. Collectively, these data prove that EtO is systemically available following inhalation exposure, and the bone marrow, the target tissue for the *Pig-a* and MN endpoints investigated in this study, was exposed to the test substance.

**Figure 2:**
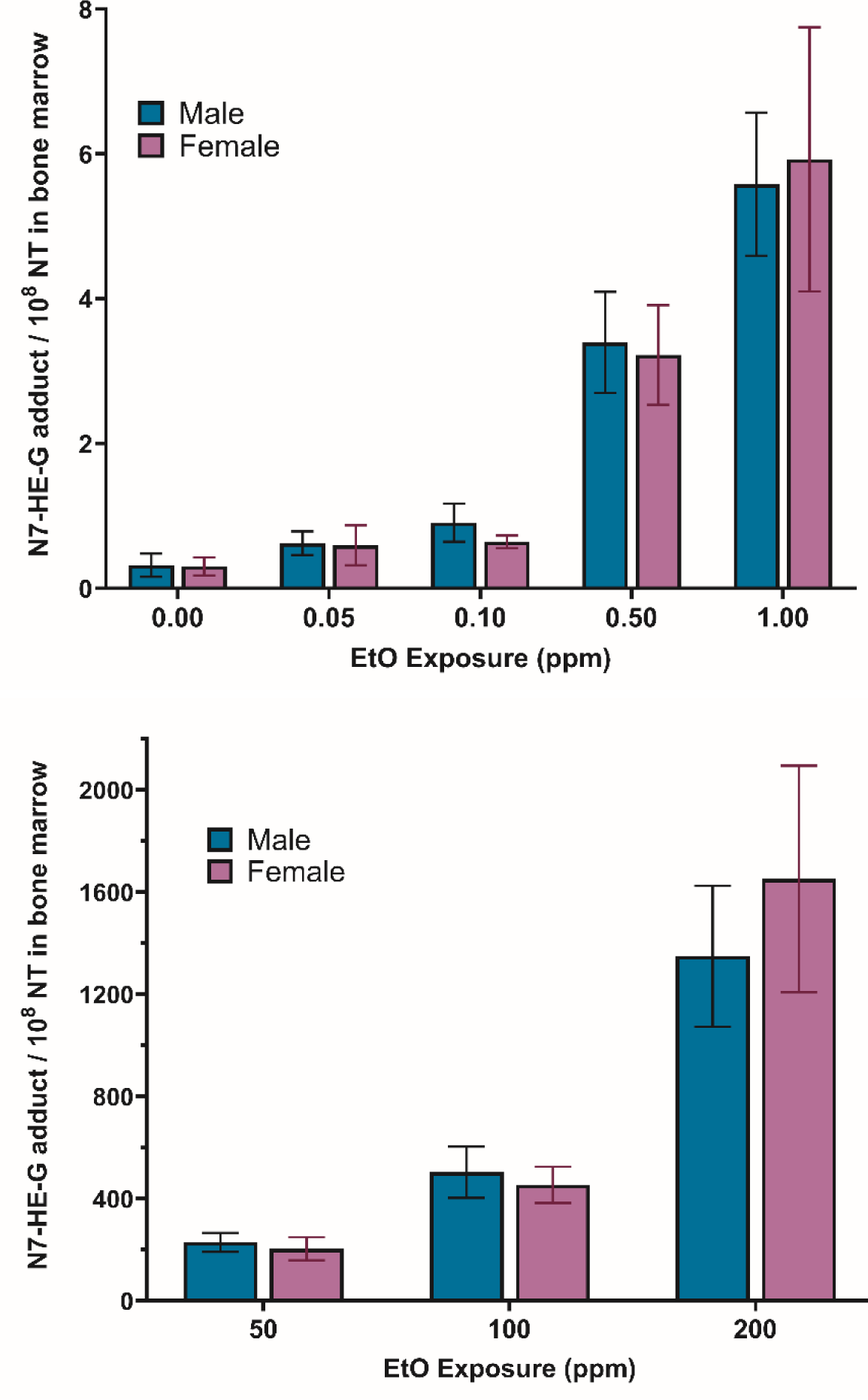
N7-HE-G Adducts per 10^8^ nucleotides (NT) in the Bone Marrow of Mice Exposed to Ethylene Oxide (EtO) for 28 Days by Whole Body Inhalation. Numerical Data (Mean ± Standard Error) reported in Chiu et al. (2026b) were plotted below in two charts to illustrate the response differences along the Y-axis at the lower (0-1 ppm) vs. higher (50-200 ppm) exposure concentrations.

The frequencies of *Pig-a* gene mutations in erythrocytes are shown in Table 1. The mutant frequencies seen in the 0-ppm group were consistent with the values reported in the literature (Olsen et al., 2017). A treatment-related and biologically significant increase in *Pig-a* mutants was seen in females exposed to 200 ppm, primarily reflecting in an increased mutant reticulocyte frequency. The adequacy of the experimental conditions used in the study for the detection of induced mutant frequencies was ascertained by the observation of markedly elevated mutant frequencies in mice dosed with the positive control chemical (ENU). These results showed a “hockey-stick” shape for the dose-response of EtO-induced *Pig-a* mutations, with no effects at lower exposures. There were significant increases in percent RET values in EtO-exposed females, with the value observed at the 200-ppm exposure suggestive of compensatory or rebound erythropoiesis in the bone marrow.

**Table 1:** Erythrocyte *Pig-a* Mutant Frequencies in B6C3F1 Mice Exposed to Ethylene Oxide(EtO) for 28 Days via Whole Body Inhalation. Values are Mean ± Standard Deviation.

| EtO Conc (ppm) | N<sup>1</sup><br>Male/Female | %RET<sup>2</sup> |  | Mutant RBC (x10<sup>-6</sup>) |  | Mutant RET (x10<sup>-6</sup>) |  |
| --- | --- | --- | --- | --- | --- | --- | --- |
|  |  | Male | Female | Male | Female | Male | Female |
| 0 (Air Control) | 30/29 | 4.00 ± 0.96 | *3.27 ± 0.56 | 0.05 ± 0.07 | *0.17 ± 0.39 | 0.34 ± 1.16 | *0.23 ± 0.34 |
| 0.05 | 10/10 | 3.94 ± 0.43 | 3.26 ± 0.33 | 0.02 ± 0.02 | 0.18 ± 0.18 | 0.13 ± 0.15 | 0.55 ± 0.62 |
| 0.1 | 10/10 | 4.16 ± 0.54 | 3.37 ± 0.50 | 0.10 ± 0.16 | 0.39 ± 0.53 | 0.04 ± 0.05 | 1.64 ± 2.34 |
| 0.5 | 10/10 | 4.27 ± 1.42 | 4.0 ± 0.79 | 0.05 ± 0.10 | 0.04 ± 0.03 | 0.68 ± 2.04 | 0.09 ± 0.13 |
| 1 | 10/10 | 5.43 ± 2.80 | <sup>3</sup>5.74 ± 3.36 | 0.03 ± 0.02 | 0.22 ± 0.42 | 0.06 ± 0.08 | 0.9 ± 1.78 |
| 50 | 20/19 | 4.02 ± 0.96 | <sup>3</sup>4.31 ± 0.74 | 0.06 ± 0.13 | 0.26 ± 0.55 | 0.35 ± 0.86 | 1.77 ± 5.00 |
| 100 | 10/10 | 4.38 ± 1.72 | <sup>3</sup>4.87 ± 1.94 | 0.02 ± 0.01 | 0.09 ± 0.16 | 0.07 ± 0.11 | 0.18 ± 0.21 |
| 200 | 10/9 | 4.64 ± 1.06 | <sup>3</sup>11.26 ± 4.75 | 0.04 ± 0.04 | <sup>3</sup>0.99 ± 0.88 | 0.10 ± 0.11 | <sup>3</sup>4.31 ± 6.62 |
| Positive Control<sup>4</sup> | 12/12 | 4.87 ± 1.49 | 5.82 ± 2.06 | <sup>5</sup>10.96 ± 4.84 | <sup>5</sup>11.02 ± 4.51 | <sup>5</sup>61.04 ± 43.22 | <sup>5</sup>38.95 ± 32.40 |

<sup>1</sup>Number of animals; <sup>2</sup>Percent reticulocytes; <sup>3</sup>Statistically significant difference(p<0.05) from the Air Control; <sup>4</sup>30 mg/kg/day of *N*-nitroso-*N*-ethyl urea administered by oral gavage on study days 1-3; <sup>5</sup>Data on mutant frequencies were judged to be significantly elevated based on marked increases vs. 0 ppm group; \*Significant trend (p<0.025).

In blood samples collected after 5 days of EtO exposure, there was a small, but statistically significant increase in MN-RET frequency at the 200-ppm exposure level (up to 40%; Table 2). A treatment-related effect on MN was not expected to manifest in NCE given the short exposure duration. The %RET values were significantly lower at the highest concentration, indicative of potential cytotoxicity, while the small, but significant, increase at the 1-ppm group was not considered to be treatment related.

**Table 2:** Erythrocyte Micronucleus (MN) Frequencies in Male B6C3F1 Mice Exposed to Ethylene Oxide (EtO) for 5 Days via Whole Body Inhalation. Values are Mean ± Standard Deviation.

| EtO Conc (ppm) | N<sup>1</sup><br>Male/Female | %RET<sup>2</sup> |  | %MN-NCE |  | %MN- RET |  |
| --- | --- | --- | --- | --- | --- | --- | --- |
|  |  | Male | Female | Male | Female | Male | Female |
| 0 (Air Control) | 30/30 | 2.21 ± 0.25 | 1.96 ± 0.45 | 0.18 ± 0.02 | *0.11 ± 0.01 | *0.27 ± 0.05 | *0.21 ± 0.03 |
| 0.05 | 10/10 | 2.30 ± 0.38 | 2.18 ± 0.51 | 0.17 ± 0.01 | 0.11 ± 0.01 | 0.30 ± 0.04 | 0.21 ± 0.02 |
| 0.1 | 10/10 | 2.32 ± 0.30 | 2.24 ± 0.37 | 0.17 ± 0.02 | 0.12 ± 0.01 | 0.26 ± 0.06 | 0.22 ± 0.04 |
| 0.5 | 10/10 | 2.34 ± 0.27 | 1.91 ± 0.50 | 0.18 ± 0.01 | 0.13 ± 0.01 | 0.29 ± 0.04 | 0.23 ± 0.05 |
| 1 | 10/10 | <sup>3</sup>2.78 ± 0.44 | 1.56 ± 0.56 | 0.18 ± 0.02 | 0.13 ± 0.01 | 0.28 ± 0.05 | 0.20 ± 0.04 |
| 50 | 20/20 | 2.11 ± 0.26 | 2.06 ± 0.41 | 0.17 ± 0.02 | 0.12 ± 0.01 | 0.30 ± 0.05 | 0.20 ± 0.03 |
| 100 | 10/10 | 2.50 ± 0.32 | 2.82 ± 2.68 | 0.19 ± 0.02 | 0.13 ± 0.01 | 0.31 ± 0.05 | 0.21 ± 0.05 |
| 200 | 10/10 | <sup>3</sup>1.79 ± 0.24 | <sup>3</sup>0.92 ± 0.41 | 0.18 ± 0.01 | 0.13 ± 0.01 | <sup>3</sup>0.35 ± 0.04 | <sup>3</sup>0.30 ± 0.08 |
| Positive Control<sup>4</sup> | 12/12 | 0.98 ± 0.28 | 0.73 ± 0.27 | 0.28 ± 0.02 | 0.20 ± 0.02 | <sup>5</sup>1.98 ± 0.55 | <sup>5</sup>1.71 ± 0.53 |

<sup>1</sup>Number of animals; <sup>2</sup>Percent reticulocytes; <sup>3</sup>Statistically significant difference (p<0.05) from the Air Control; <sup>4</sup>30 mg/kg/day of *N*-nitroso-*N*-ethyl urea administered by oral gavage on study days 1-3; <sup>5</sup>Data on mutant frequencies were judged to be significantly elevated based on marked increases vs. 0 ppm group; \*Significant trend (p<0.025).

Like the 5-day samples, a small (up to 44%), but statistically significant increase in MN-RET frequency was also observed in blood samples collected on Study Day 28 at concentrations of 1-200 ppm (Table 3). The MN-NCE values in females exposed to 1 to 200 ppm and males in the 200-ppm group were identified as significantly different from the corresponding 0-ppm controls. The increases seen in females exposed to 1 to 100 ppm need to be viewed with caution about their biological significance given their minor differences from the controls and the absence of a significant effect on the leading indicator (i.e., MN-RET) of an induced effect. However, it is acknowledged that such a correspondence is not mandatory since MN-NCEs represent the cumulative effect over the 28 days of exposure, while MN-RETs represent recently induced damage, and the MN-RET rapidly (within 2 days) mature into NCE after entering the blood circulation. The %RET values of the 200-ppm group were significantly lower than the air control group, suggesting a potential cytotoxic effect of the treatment. Mice treated with the positive control chemical had up to a 700% increase in the incidence in MN-RET, confirming the adequacy of the experimental conditions to detect an induced effect.

**Table 3:** Erythrocyte Micronucleus (MN) Frequencies in B6C3F1 Mice Exposed to Ethylene Oxide for 28 Days via Whole Body Inhalation. Values are Mean ± Standard Deviation.

| Conc (ppm) | N<sup>1</sup><br>Male/Female | %RET<sup>2</sup> |  | %MN-NCE |  | %MN- RET |  |
| --- | --- | --- | --- | --- | --- | --- | --- |
|  |  | Male | Female | Male | Female | Male | Female |
| 0 (Air Control) | 30/30 | 1.22 ± 0.11 | 0.84 ± 0.17 | *0.19 ± 0.02 | *0.11 ± 0.01 | *0.27 ± 0.05 | *0.17 ± 0.03 |
| 0.05 | 10/10 | 1.26 ± 0.18 | 0.93 ± 0.12 | 0.19 ± 0.02 | 0.10 ± 0.01 | 0.29 ± 0.03 | 0.16 ± 0.03 |
| 0.1 | 10/10 | 1.41 ± 0.17 | 0.88 ± 0.22 | 0.18 ± 0.02 | 0.11 ± 0.00 | 0.28 ± 0.05 | 0.17 ± 0.02 |
| 0.5 | 10/10 | 1.20 ± 0.12 | 0.97 ± 0.18 | 0.19 ± 0.02 | 0.11 ± 0.01 | 0.26 ± 0.03 | 0.17 ± 0.03 |
| 1 | 10 /9 | 1.22 ± 0.13 | 0.91 ± 0.20 | 0.18 ± 0.02 | <sup>3</sup>0.12 ± 0.01 | 0.28 ± 0.03 | 0.20 ± 0.02 |
| 50 | 20/20 | 1.30 ± 0.11 | 1.07 ± 0.17 | 0.19 ± 0.02 | <sup>3</sup>0.12 ± 0.01 | 0.28 ± 0.03 | 0.19 ± 0.04 |
| 100 | 10/10 | 1.33 ± 0.09 | 0.82 ± 0.15 | 0.20 ± 0.02 | <sup>3</sup>0.13 ± 0.01 | 0.30 ± 0.04 | 0.19 ± 0.02 |
| 200 | 10/10 | <sup>3</sup>1.77 ± 0.60 | 2.13 ± 0.84 | <sup>3</sup>0.23 ± 0.01 | <sup>3</sup>0.19 ± 0.03 | <sup>3</sup>0.34 ± 0.02 | <sup>3</sup>0.23 ± 0.06 |
| Positive Control<sup>4</sup> | 12/12 | 0.40 ± 0.16 | 0.50 ± 0.19 | 0.19 ± 0.02 | 0.12 ± 0.01 | <sup>4,5</sup>1.80 ± 0.22 | <sup>4</sup>1.68 ± 0.26 |

<sup>1</sup>Number of animals; <sup>2</sup>Percent reticulocytes; <sup>4</sup>25 mg/kg/day of cyclophosphamide monohydrate administered by oral gavage on study days 25 and 26; <sup>3</sup>Statistically significant difference (p<0.05) from the Air Control; <sup>4</sup>Data on MN frequencies were judged to be significantly elevated based on marked increases vs. 0 ppm group rather than statistical analyses; <sup>5</sup>One MN-RET value was not calculated because %RET was below 0.1%, so this calculation was based on N = 11 rather than 12; \*Significant trend (p<0.025).

The dose-response for the induction of *Pig-a* mutations and MN is graphically shown in Figures 3 and 4, respectively. Data for only the 28-day females are shown because they are representative of males, as well as the 5-day time point for MN. There was no evidence for the hypothesized 2-piece linear spline response for these endpoints across the concentrations from 0.05 to 200 ppm.

**Figure 3:**
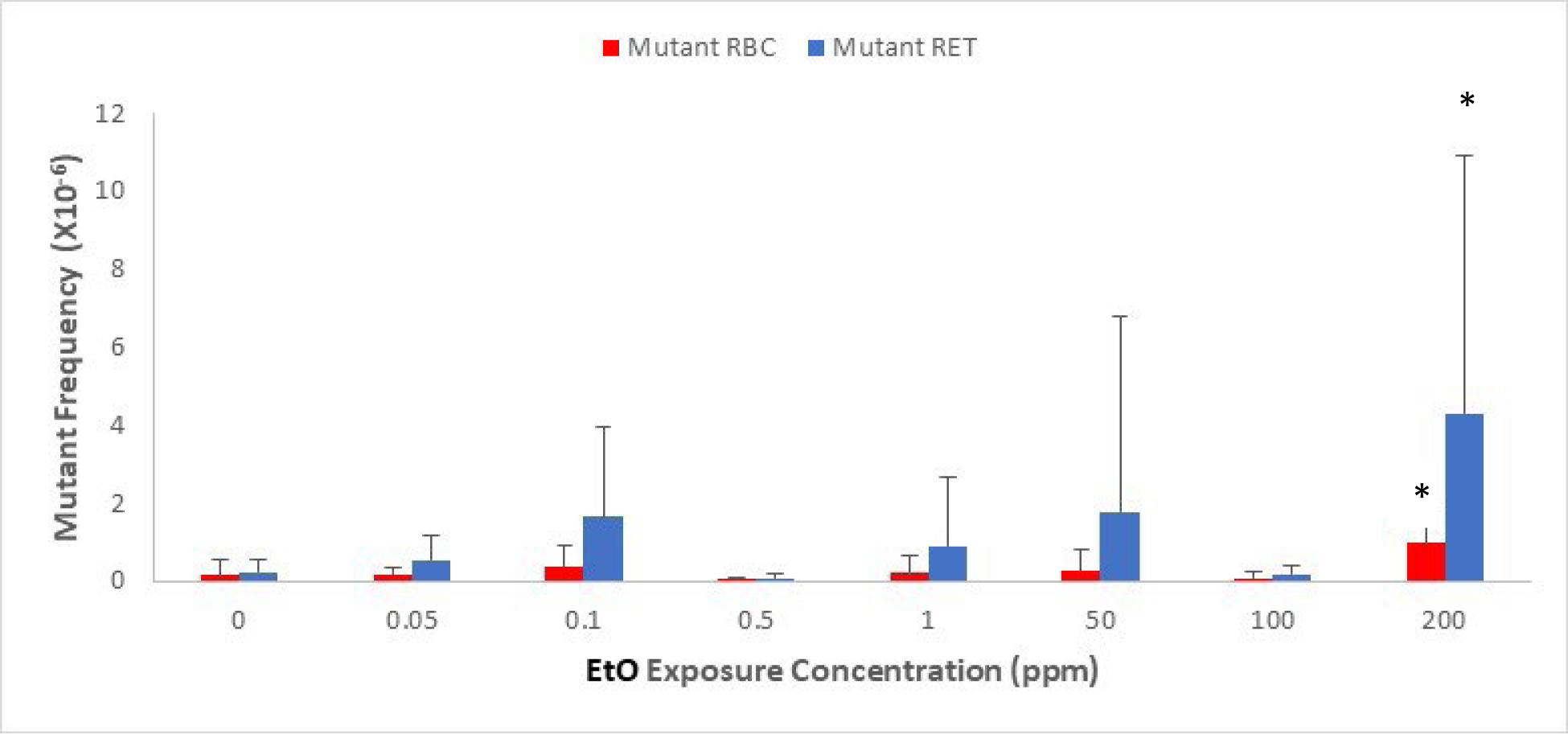
Dose-Response for *Pig-a* Mutation Frequencies in Female Mice Exposed for 28 Days to Ethylene Oxide by Whole Body Inhalation. *Significantly different from 0 ppm.

**Figure 4:**
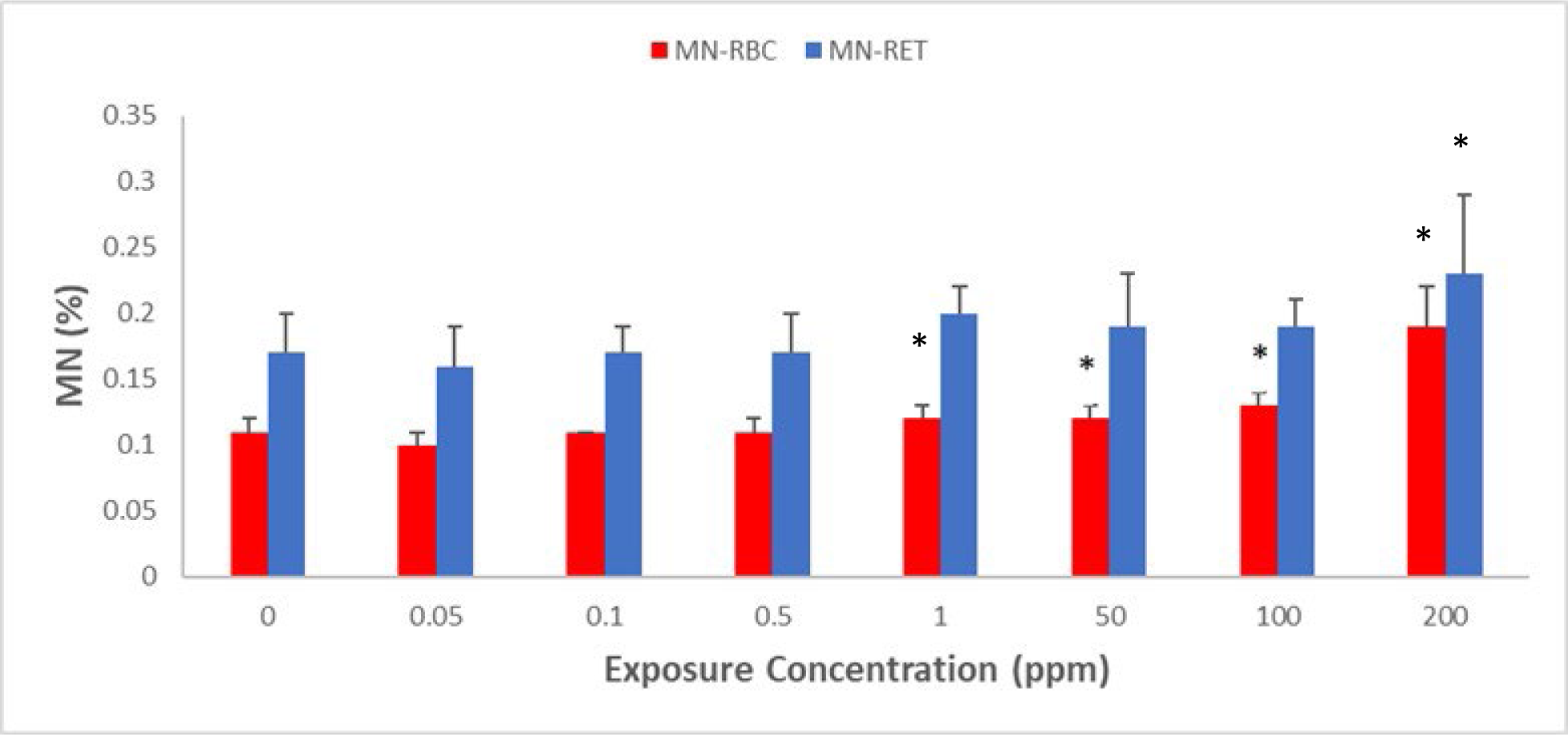
Dose-Response for Micronucleus Induction in Female Mice following 28 Days of Whole-Body Inhalation Exposure to Ethylene Oxide. *Significantly (p<0.05) different from 0 ppm.

## DISCUSSION

The findings of this study are significant in that they provide a biologically plausible basis that is based on the presumed apical MOA key events (i.e., mutation and cytogenetic damage) for the selection of a statistical dose-response model for EtO cancer risk assessment. During quantitative cancer risk assessment, the observed data can be fit, with equal statistical validity, using different dose-response models, resulting in vastly different risk estimates. This is the case with EtO cancer risk assessment using lymphoid cancer mortality data from epidemiology studies, where the 1 in a million-cancer risk derived by the US EPA using the 2-piece linear spline model (i.e., 0.0001 ppb) is nearly three orders of magnitude lower than that estimated by TCEQ using the established and conservative log-linear CPH model (i.e., 0.24 ppb) for linear no-threshold approach. The TCEQ model is thus linear throughout the entire range of cumulative exposure for lymphoid cases, which is consistent with the biological evidence based on the early key event (i.e., mutagenicity) of the mutagenic cancer MOA for EtO. The 2-piece linear spline model used by the US EPA, on the other hand, has two linear CPH models connected by a knot at 1600 ppm-days that was determined statistically based on the local maximum likelihood. The first linear CPH model has a very steep slope, and the second one has a shallow slope. The *in vivo* dose-response data for the genotoxicity endpoints observed in this study do not provide support for this model selection, especially given the current and previous data indicating EtO is a relatively weak mutagen in that it requires high exposure concentration and long exposure durations to elicit its mutagenic response.

The molecular initiating event for EtO-induced mutagenicity is its reactivity with the nucleophilic centers of DNA. The dose-response seen for the induction of DNA adducts (N7-HE-G and O^6^-HE-dG) from the animals exposed in this study was reported by Liu et al. (2026b). Their analysis showed an exposure-dependent increase in the non-mutagenic N7-HE-G adducts at all concentrations of EtO tested. However, the pro-mutagenic O^6^-HE-dG adducts were detected only at exposures ≥50 ppm. The absence of O^6^-HE-dG at lower exposures is not due to a lack of analytical sensitivity since the study used an extremely sensitive analytical method for its detection, i.e., a detection limit of 3 amol equivalent to detection of less than one O^6^-HE-dG adduct/cell. Of particular importance, the dose-response slopes for both adducts increased disproportionately only at higher concentrations of EtO (i.e., ≥50 ppm) presumably after all the detoxification and repair capacities were saturated. Importantly, the most abundant N7-HE-G adduct being a biomarker of exposure of the critical target (i.e., DNA) showed no indication of a dose response pattern reflecting an initial steeper linear slope with subsequent plateauing at higher concentration. In fact, this adduct showed a contrasting dose-response pattern that transitioned to a steeper slope only at higher exposure concentrations. The disproportionate increase in the adduct load at higher EtO exposure concentrations was consistent with the disproportionate increase in blood concentrations as predicted by the PB-PK model of Fennel and Brown (2001)), and the upward bend in blood concentrations was attributable to concurrent dose-dependent depletion of glutathione essential for conjugation-mediated detoxification at the higher EtO exposures. The upward bend in blood concentrations was attributable to saturation of glutathione conjugation-mediated detoxification at the higher EtO exposures. Overall, these data provide further biological evidence in support of the linear CPH model as preferred and conservative approach for risk assessment of EtO carcinogenicity and shows the biological implausibility of the 2-piece linear spline dose-response model.

It is worth considering whether the findings from this study, which focused on a 28-day exposure period, are relevant to longer-term exposures seen in animal cancer bioassays or in real-life human situations. As stated earlier, the 28-day exposure period is adequate to reach steady state for the induction of DNA adducts in the mouse, rat and humans (Filser and Klein, 2017; Walker et al., 2000). Adduct loads at steady state are influenced by their formation, DNA repair, loss and cell turn over. Accordingly, exposures longer than 28 days are not expected to yield results different from those derived in the current study. Importantly, it needs to be pointed out that the apical effects for gene mutations and chromosomal damage were seen primarily at the 200-ppm exposure. Despite this, the dose-response for these effects was conservatively assumed to be linear with no thresholds as a worst-case scenario for this DNA-reactive chemical. Thus, it is extremely implausible that longer exposure durations would somehow induce effects with a dose-response pattern like the one predicted by the 2-piece linear spline model used by the US EPA (2016) for EtO cancer risk assessment (Gollapudi et al., 2020). Additionally, for chemicals acting through a genotic MoA, mutation induction is an early initiating event in the cancer pathway rather than appearing late during the chronic exposure (Moore et al., 2008).

An important question in the context of this study is the relevance, or lack thereof, of the data from mice to select a model for human cancer risk assessment, especially when the cancer data used for risk assessment were from human epidemiology studies. In general, DNA repair capacity of humans is similar, if not better, than that of mice (MacRae et al., 2015). This also applies to *O*^6^-methylguanine-DNA methyltransferase (MGMT), a highly conserved repair protein involved in the direct repair pathway of the pro-mutagenic O^6^-HE-dG and other *O*^6^-dG alkyl adducts (Roy et al., 1995; Klapacz et al., 2016). In addition, EtO PB-PK modeling indicates that systemic blood concentrations are similar between mice and humans at EtO exposures ≤100 ppm, suggesting that EtO detoxification toxicokinetics operate similarly across the species (Fennell and Brown, 2001). Thus, the mouse is conservatively a valid surrogate model to predict the genotoxicity of EtO in humans, especially at exposure levels that do not overwhelm the detoxification pathways critical to maintaining homeostasis. At exposure levels >200 ppm, PB-PK modeling indicates that humans are better equipped to maintain EtO detoxication (e.g., through glutathione conjugation) and thus the response in the mouse can again be considered as a worst-case scenario (Fennel and Brown, 2001). Collectively, the overall MOA data derived from mice in this study and others conservatively justify selection of a single-linear (CPH) dose response model for the risk assessment EtO carcinogenicity.

Results from this study, informing the weak *in vivo* mutagenicity of EtO, are consistent with previously reported studies in the bone marrow (Recio et al., 2004) and the lung (Manjanatha et al., 2017) tissues of Big Blue^®^ transgenic mice. In both cases, the shape of the dose response is at best conservatively characterized as having a single linear slope. In fact, significant increases in mutations were only observed at the higher doses of EtO used in these studies (i.e., ≥100 ppm in the bone marrow and at 200 ppm in the lung), and that too after extended durations of exposure (i.e., after 48 weeks but not at 12 or 24 weeks in the bone marrow and at 8, but not at 4 weeks in the lung). In a study reported only as a meeting abstract, LeBaron et al. (2013) did not see MN induction in erythrocytes of B6C3F1 mice exposed for 28 days to EtO concentrations ranging from 10-200 ppm. The relatively weak mutagenicity of EtO was also noted by Gollapudi et al. (2020) in their review of 40 *in vivo* data sets to derive benchmark dose levels for gene mutations and chromosomal damage.

A steeper linear slope at lower doses followed by a shallower slope at high doses for the mutagenicity of EtO, as predicted by the 2-piece linear spline model, is also not conceivable on mechanistic considerations. First, EtO is a direct-acting mutagen that does not require metabolic activation for its DNA reactivity. Thus, potential saturation of metabolic activation at higher concentrations leading to a shallower slope, e.g., saturation of vinyl chloride metabolism to its carcinogenic epoxide (Slikker et al., 2004), is not expected for the mutagenicity of EtO. Secondly, the highest exposure level evaluated in the current study, i.e., 200 ppm, is twice the tumorigenic level in the carcinogenicity study using the same strain of mice (NTP, 1987). There was no evidence for excessive cytotoxicity even at this concentration, which shows that attenuation or even a decrease in the manifestation of genotoxicity at higher doses is not a biological possibility. This, in conjunction with no detectable genotoxicity at the lower doses possibly related to efficient detoxification and/or DNA repair, makes the 2-piece linear spline response biologically implausible for this key event (i.e., genotoxicity).

Dose response studies interrogating mutagenicity in cancer driver genes might be more informative than those using reporter endpoints such as the *Pig-a* gene because of their causal association with carcinogenicity. Toward this aim, Parsons et al. (2013) investigated the induction of mutations at codon 12 of the *KRas* gene in lung tissues of mice exposed to EtO for various concentrations and durations. Mutations at this codon have been implicated in the etiology of mouse lung tumors and specifically, all mouse lung tumors in the NTP inhalation study on EtO have the *Kras* mutations at this codon (Hong et al., 2007). Parsons et al. (2013) observed that following 4 weeks of EtO exposure, but not at 8 or 12 weeks, *KRas* mutant fraction at codon 12 as a function of cumulative EtO dose increased in a non-monotonic manner, and the observed response was consistent with the amplification of preexisting mutations, rather than through genotoxicity. Clonal amplification of pre-existing mutations is usually the result of cell proliferation triggered by either mitogenicity or cytotoxicity of the treatment, a MOA inconsistent with the mutagenic MOA action conservatively envisaged for EtO-induced tumors.

## CONCLUSION

The dose response pattern for genetic damage, which is the first key event of the mutagenic MOA for the carcinogenicity of EtO, supports the biological plausibility of a single-slope linear CPH, rather than the 2-piece spline, model for the derivation of inhalation unit cancer risk for this chemical.

## Statement of Author Contribution

BBG, AL, JEB and KL participated in designing the study. PC and JTW were involved in conducting the in-life phase of experimental work. JCB, DT, and SD collected and analyzed the flow-cytometric data. KL provided DNA adduct data. BBG prepared the first draft of the manuscript. All authors critically reviewed the draft and approved the final version for publication.

## Acknowledgements

Technical discussions with toxicologists of the sponsoring industry consortium are duly acknowledged. The authors also thank the contributions of various laboratory personnel at Charles River Laboratories, Ashland, OH (USA) and Ms. Svetlana Avlasevich of Litron Laboratories, Rochester, NY (USA) for their contributions in conducting this complex study.

## Data Availability

Reasonable requests for the original data can be made to the American Chemistry Council Ethylene Oxide Panel, 700 2nd Street NE, Washington, DC 20002, USA.

## Footnotes

1 Due to technical issues, 0.05 and 0.1 ppm groups were exposed for 27 days in Phase I.

